# A non-invasive method for profiling the gut microbiome and virome of honey bee queens

**DOI:** 10.1101/2025.04.15.648983

**Authors:** L.A. Holmes, P. Wolf Veiga, J.S. Pettis, M.M. Guarna, S.E. Hoover

## Abstract

High honey bee colony mortality worldwide has underscored the critical role of queen bee health in colony survival, with poor queen quality frequently linked to colony losses. The gut microbiome plays fundamental roles in immunity, nutrition, and reproduction, making its characterization essential for understanding stressors that impact queen health, longevity, and fecundity, yet its role in mediating stress responses remains poorly understood. Here, we present a novel, non-invasive method for collecting feces from queen honey bees and demonstrate its potential as a powerful tool for profiling the gut microbiome, detecting stressor exposure, and screening for viral infections. This approach permits repeated, longitudinal assessments of individual queens, providing unprecedented insights into how environmental and pathogenic pressures influence queen health, longevity, and reproductive capacity. Beyond research applications, benefits include evaluating queens before colony introduction and mitigating disease transmission risks in international trade, where pathogen spread remains a major regulatory challenge.

---

Next generation sequencing enables researchers to identify microorganisms associated with animal hosts without the need for cultivation, significantly advancing the study of gut microbiome ecology and host interactions (Maritan *et al.,* 2024; Peixoto, Harkins, and Nelson, 2021). However, generalizing the effects of the gut microbiome on hosts is challenging due to the immense variation in gut microbiomes across different animal taxa (as recently reviewed in Maritan *et al.,* 2024). Despite this complexity, clear host associations and critical roles of the gut microbiome in host immunity, nutrition, and reproduction have been amply demonstrated for both vertebrate (as reviewed in Peixoto, Harkins, and Nelson, 2021; and Hooper *et al*., 2012) and invertebrate animals (Motta and Moran, 2024; Douglas, 2019; and Engel and Moran, 2013). In vertebrates, the gut microbiome is often studied using fecal samples, which provide an accurate proxy of the gut microbial composition (Yan *et al.,* 2019; Härer and Rennison, 2023) However, differences between fecal and whole-gut sequencing have been noted (see Ahn *et al.,* 2023). In contrast, studying the gut microbiome of invertebrates has typically required destructive sampling of the entire host specimen, as individual fecal samples are typically too small for DNA extraction and sequencing (but see Franz *et al.,* 2023; Moeller *et al.,* 2020; O’Hara *et al.,* 2020; Zhu *et al.,* 2019; Fink *et al.,* 2013 for fecal collections pooled from multiple individuals).

The honey bee (*Apis mellifera*) is a valuable model for studying the gut microbiome due to its well-documented biology and critical role in agriculture. The honey bee gut microbiome is relatively simple (Kwong and Moran, 2016), yet highly specialized and conserved, to the extent that 95% of the honey bee gut community is dominated by nine bacterial species, of which five are core species found in honey bee workers throughout the world (reviewed in Kwong and Moran, 2024; 2016 and Zheng *et al.,* 2018). This conserved gut microbiome plays a crucial role in honey bee health and disease resistance (reviewed by Motta and Moran, 2024). For example, honey bees harbouring a normal gut microbiota show increased expression of antimicrobial peptides essential for immunity (Kwong, Mancenido, and Moran, 2017). *In situ* studies further demonstrate that opportunistic pathogens persist when core honey bee gut microbiota are absent but are rapidly eliminated in their presence (Steele *et al*., 2021). For example, core gut symbionts such as *Snodgrassella alvi*, help induce antimicrobial peptide production, protecting honey bees from opportunistic pathogens (Horak, Leonard, and Moran, 2020).

Queen honey bees are typically the only reproductive female in the colony, making their health a top priority for apicultural industries worldwide. Although research on the queen gut microbiome is limited, studies indicate that it is highly structured and consists of a few core bacterial groups, including Firmicutes (e.g., *Bombilactobacillus* (formerly *Lactobacillus* Firm4) and *Lactobacillus*), Alphaproteobacteria (e.g., *Bombella*), and Actinobacteriota (e.g., *Bifidobacterium*) (Copeland, Anderson, and Mott, 2022). However, the queen’s gut microbiome only partially overlaps with those of worker bees. Compared to workers, queens have a less diverse bacterial community (Powell *et al*., 2018), with Firmicutes and Alphaproteobacteria as the dominant groups (Copeland, Anderson, and Mott, 2022; Wang *et al*., 2022; Powell *et al*., 2018; Tarpy *et al*., 2015). Additionally, the composition of the queen gut microbiome varies with age (Tarpy *et al*., 2015) and environmental factors (Copeland *et al*., 2024).

The relatively simple gut microbiome of honey bees provides a valuable model for studying how specialized gut microbiome communities assemble and interact with their hosts. Understanding these interactions is essential for identifying stressors that impact queen health and reproduction. Beekeepers frequently cite ‘queen failure’ as a cause of honey bee colony mortality during winter (Claing *et al*., 2023), with many reporting the need to replace half their queens annually (Amiri *et al*., 2017). However, the causes of queen failure are often unclear but may result from poor mating (Pettis *et al*., 2016), exposure to pesticides (Walsh *et al*., 2020; DeGrandi-Hoffman, Chen, and Simonds, 2013; Collins *et al.,* 2004), pathogens (Amiri *et al.,* 2020; DeGrandi-Hoffman, Chen, and Simonds, 2013; Gauthier *et al.,* 2011), temperature stress (McAfee *et al.,* 2020), poor nutrition (Fine *et al.,* 2018), or a combination of these factors.

The health of new queens purchased from queen production companies is often uncertain, especially in temperate regions that rely on spring imports of queens from warmer regions (AAFC, 2022; Bixby *et al*, 2020; 2019). During shipment and transit, queens may experience additional stress, including exposure to extreme temperatures (Pettis *et al*., 2016). Previous studies have shown that transported queens can be exposed to temperatures between 4°C and 38°C (Holmes *et al*., 2023; McAfee *et al*., 2020; Pettis *et al.,* 2016), which can negatively affect their reproductive viability. Spermatozoa stored in the queen’s spermatheca (i.e., an organ the queen uses to store sperm after mating to use for the remainder of her life) are particularly sensitive to temperature fluctuations. For example, exposure to temperatures below 10°C and above 40°C can significantly reduce sperm viability (McAfee *et al*., 2020; Rousseau *et al*., 2020; Pettis *et al*., 2016) due to the high sensitivity of sperm cells to thermal stress (Sales *et al*., 2018; Pettis *et al*., 2016; Stürup *et al*., 2013). In addition, previous research has shown a trade-off between sperm storage and immune function in queen honey bees; investing in the long-term sperm preservation comes at a cost to immune defenses (McAfee *et al*., 2021).

Temperature stress is also known to disrupt insect microbial communities (reviewed in Mason and Shikano, 2023), however, its impact on the gut microbiome in queen honey bees remains unexplored. It is unclear whether temperature affects queen fertility (McAfee *et al*., 2020; Rousseau *et al*., 2020; Pettis *et al*., 2016) through direct effects on spermatozoa, shifts in the bacterial communities in response to temperature, or both. Currently, assessing key queen quality traits including the microbiome, proteome, pathogen load, and sperm quality requires destructive sampling, necessitating the sacrifice of the queens (Amiri *et al.,* 2020; McAfee *et al.,* 2020; Pettis *et al.,* 2016). As a result, these traits can only be evaluated retrospectively, after exposure to stressors, without any *a priori* knowledge of the queen’s original health and fertility. Furthermore, queens subjected to destructive sampling cannot subsequently go on to lead a colony.

A major challenge in the beekeeping industry is therefore the lack of non-destructive tools to assess queen honey bee health. While fecal samples have been used to study gut microbiomes in other bee species (Hotchkiss, Forrest, and Poulain, 2024; Näpflin and Schmid-Hempel, 2018), their use in honey bees has not been previously reported, and characterizing the fecal microbiome of individual bees remains limited. To address this challenge, we developed and validated a novel non-destructive method to assess queen health through the characterization of their gut microbiome and virome through fecal sampling. Here, we present a detailed protocol for collecting feces from queens and outline five key objectives: 1) determine the minimum number of fecal deposits required for successful 16S rRNA amplicon sequencing, 2) evaluate whether queen feces provide a reliable proxy for gut microbiome characterization; 3) explore the potential for fecal analyses to indicate exposure to temperature stress, 4) refine collection methods to improve DNA quality, and 5) assess the feasibility of using queen feces for virus screening. This approach provides a groundbreaking opportunity to repeatedly monitor the gut microbiome and virome of individual queens throughout their adult lifespan, assess their response to environmental stressors, and evaluate their health prior to importation and colony installation.

## RESULTS

### 1) Determining the number of fecal samples needed for successful 16S rRNA amplicon sequencing

Our sequencing results from 21 fecal and 21 gut samples returned 1,083,967 reads among 87 ASVs. After FilterandTrim (DADA2), 89.92% of the sequencing reads were retained. Sequence data from the pooled fecal deposit samples (N = 21) were rarefied to a sequencing depth of 1500 reads, where our final dataset included 337,231 sequence reads, or 98.21% of the total filtered reads from the 21 fecal samples.

The best model for predicting the number of 16S amplicon sequence reads included the number of fecal deposits pooled per sample (Figure 1, Table S2), where samples with a single fecal deposit had significantly fewer sequence reads than samples with two or more pooled fecal deposits. However, the number of sequence reads did not differ among samples with two, three, or four pooled fecal deposits, indicating at least two fecal deposits should be pooled for 16S amplicon sequencing (Figure 1).

**Figure 1.**
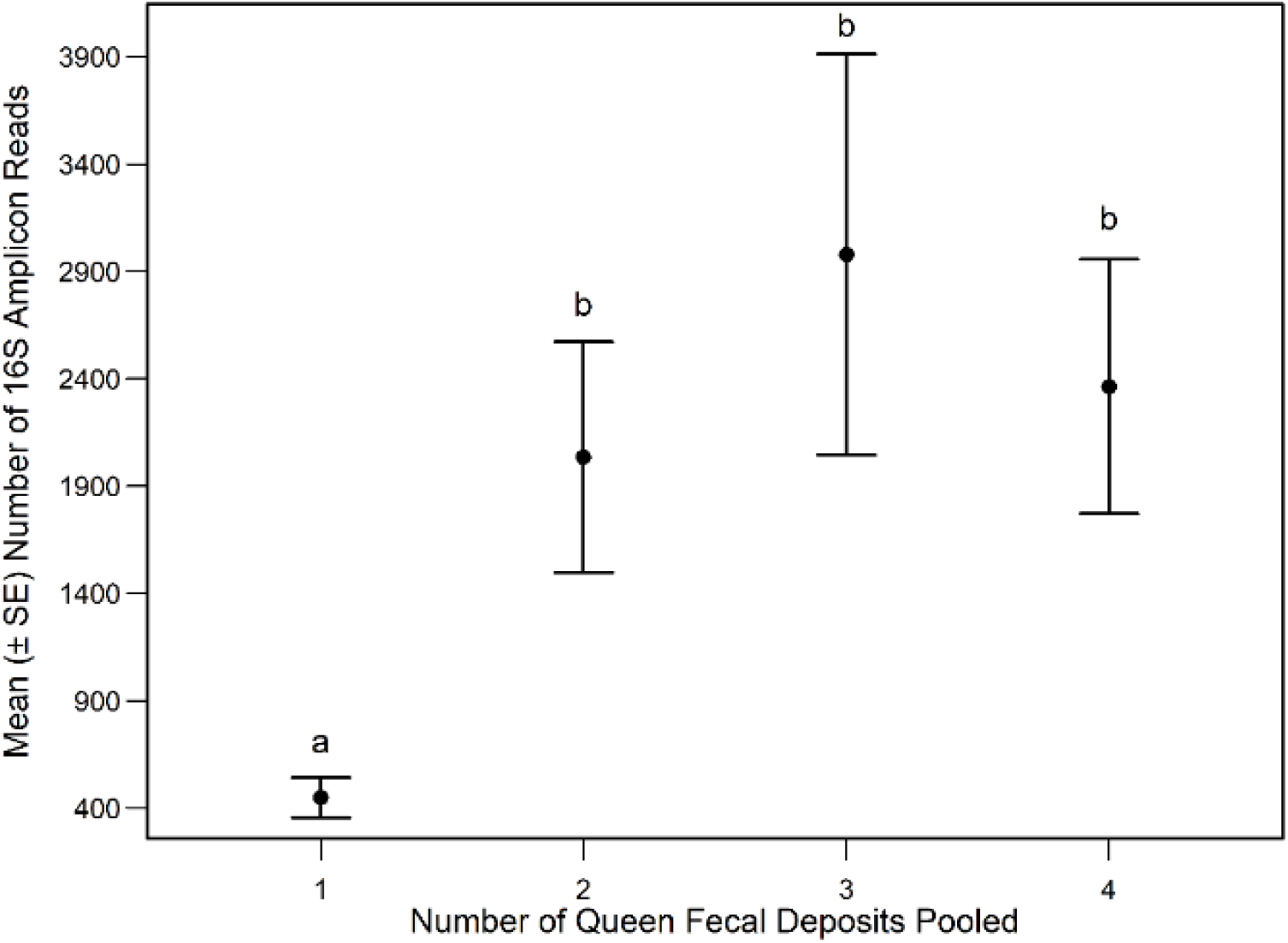
Mean (± SE) number of 16S amplicon sequence reads across the number of queen honey bee fecal deposits pooled in a sample (i.e., 1, 2, 3, or 4 pooled fecal deposits per sample; Table 1). Data were rarefied to a sequencing depth of 1500 reads. Significantly different means and standard errors of 16S amplicon sequence reads across the number of fecal deposits pooled are indicated by different letters (P < 0.05) after performing general linear hypothesis post-hoc testing on the top model selected by Akaike Information Criterion (Table S2).

**Table 1.** Queen honey bee fecal (n = 21) and gut (n = 21) samples pooled separately by queen. Queens providing the fecal deposits in a pooled fecal sample are the same queens as the pooled gut samples. A subset of the queens (N = 31) was divided into heat stress (i.e., 42°C) and control (i.e., 32°C) temperature treatments to address our third objective. The remaining queens (N = 16) were sampled from their colonies without any experimental treatment.

| Experimental Treatment (if applicable) | No. of Pooled Fecal Deposits | No. of Pooled Queen Guts |
| --- | --- | --- |
| Heat Stress | 1 | 4 |
| Heat Stress | 4 | 2 |
| Heat Stress | 2 | 1 |
| Heat Stress | 2 | 3 |
| Heat Stress | 3 | 3 |
| Heat Stress | 1 | 1 |
| Control | 1 | 4 |
| Control | 1 | 1 |
| Control | 1 | 2 |
| Control | 2 | 3 |
| Control | 4 | 1 |
| Control | 4 | 2 |
| Control | 1 | 1 |
| Control | 3 | 3 |
| No Treatment | 1 | 1 |
| No Treatment | 3 | 2 |
| No Treatment | 3 | 3 |
| No Treatment | 4 | 4 |
| No Treatment | 2 | 2 |
| No Treatment | 4 | 3 |
| No Treatment | 1 | 1 |

### 2) Characterizing the fecal and gut microbiome communities of queen honey bees

Our sequencing results from 19 fecal and 19 gut samples returned 1,082,671 reads among 87 ASVs. After FilterandTrim (DADA2), 89.94% of the sequence reads were retained in the 19 fecal and 19 gut samples. Sequence data were rarified to a sequencing depth of 2500 reads, and our dataset included 821,377 sequence reads for analyses, or 95% of the total filtered reads. Rarifying to a sequencing depth of 2500 reads removed two ASVs that were each only found in two separate paired fecal and gut sample and thus, could not be used in our fecal and gut microbiome correlation analyses.

Seven bacterial genera including *Bombilactobacillus (previously Lactobacillus* Firm-4), *Lactobacillus, Apilactobacillus* (Firm-5), *Bombella* and *Commensalibacter* (Alphaproteobacteria), *Snodgrassella* (Gammaproteobacteria), and *Bifidobacterium* (Actinobacteria) comprised 95% of the filtered and rarified reads from the 19 fecal and 19 gut samples. We found significant positive correlations between the log abundance of 16S amplicon sequence reads found in queen feces and queen gut tissues for each bacterial genera (Figure 2, Table S3), suggesting the fecal microbiome is representative of the gut microbiome found in queen honey bees. In addition, the analysis of similarities among the paired 19 fecal and 19 gut tissue samples was not significant (ANOSIM R = 0.04, p = 0.15), suggesting the bacterial communities sequenced from the queen honey bee fecal samples are similar to their paired queen gut samples (Figure 3).

**Figure 2.**
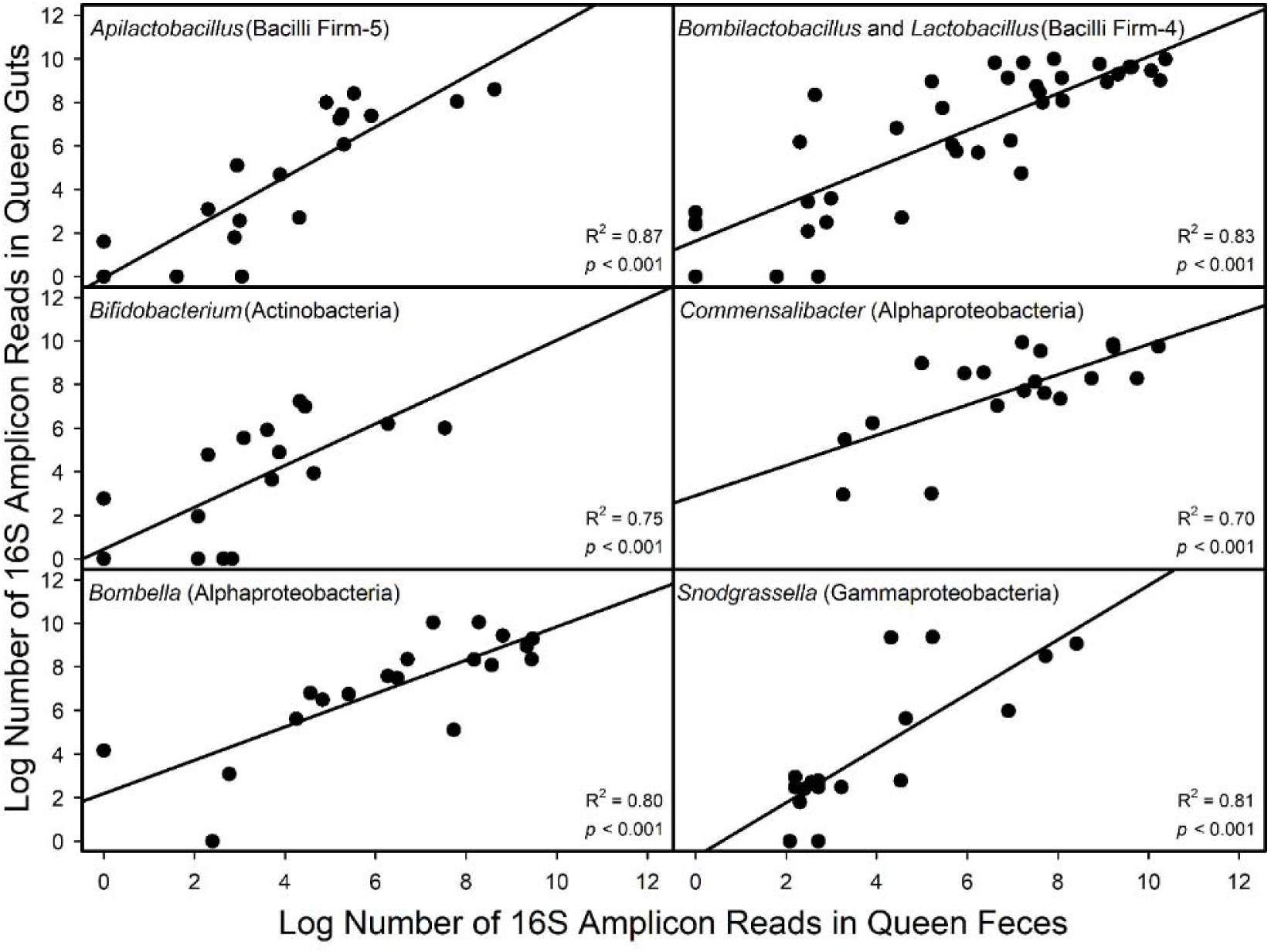
Linear relationships between the log abundance of 16S amplicon sequence reads found in paired queen gut tissues (y-axis) and fecal deposits (x-axis) for each bacterial genera represented in 95% of the filtered and rarified sequence reads from 19 fecal and 19 gut tissue samples. Pearson correlation coefficients (R^2^) and p-values are indicated for each bacterial genera Pearson correlation. Bacterial classes are indicated in parentheses.

**Figure 3.**
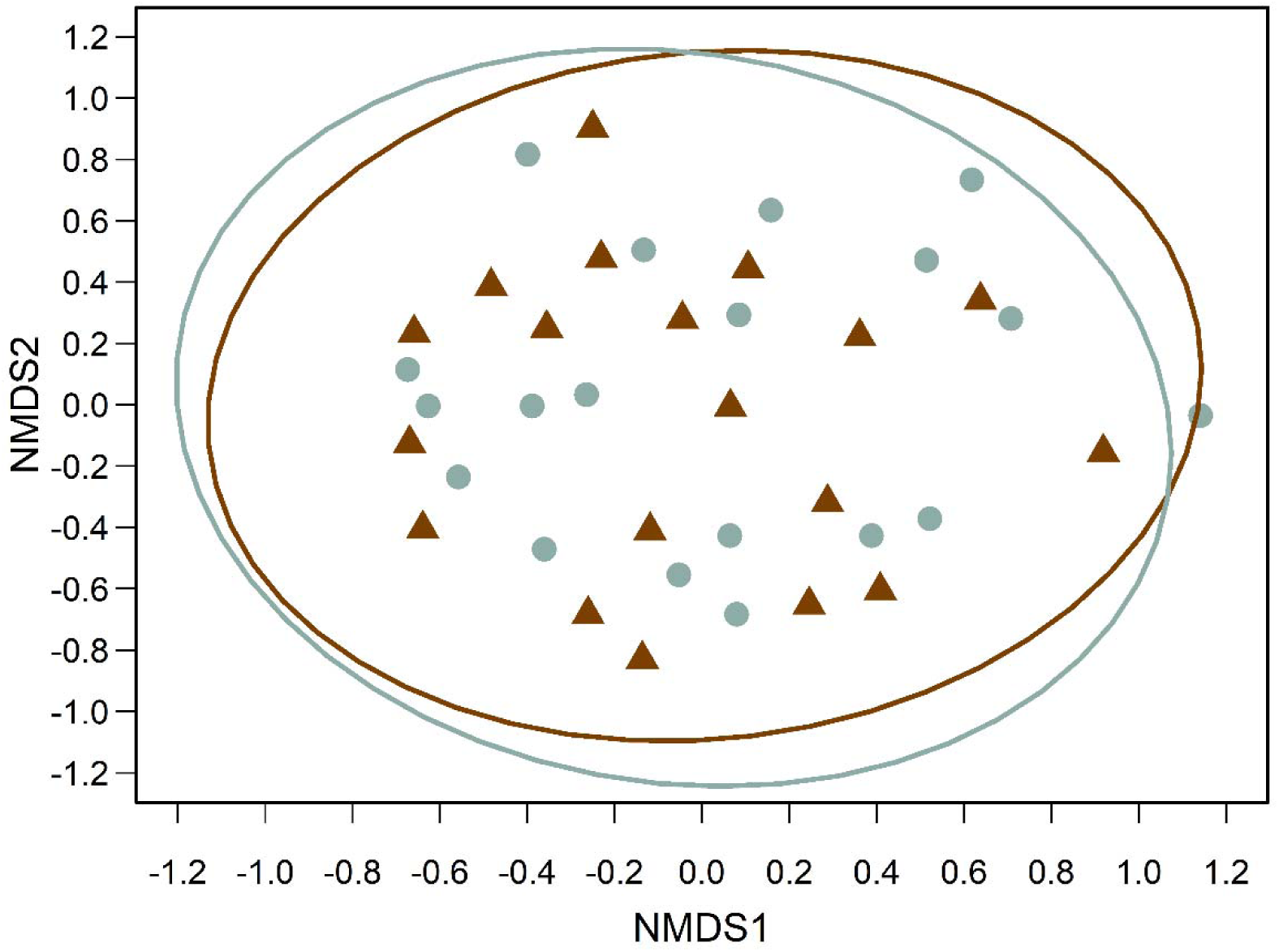
Non-metric multidimensional scaling (NMDS) plot based on a pairwise Bray-Curtis dissimilarity matrix of the bacterial community composition among paired queen honey bee fecal (i.e., triangles) and gut (i.e., circles) samples sequenced using 16S rRNA amplicon sequencing. Ellipses represent the 95% confidence intervals for fecal and gut samples in brown and gray, respectively.

### 3) Fecal and gut microbiome communities of queen honey bees in response to temperature stress

Our sequencing results from 19 fecal and 21 gut samples returned 1,083,967 reads among 87 ASVs. After FilterandTrim (DADA2), 89.83% of the sequence reads were retained in the 19 fecal and 21 gut samples. Pooled fecal deposit (n = 12) and pooled gut (n = 14) samples that were from queens assigned either heat stress or control temperature exposure treatments (Table 1) included 523,475 sequence reads for analyses after rarifying to a sequencing depth of 2000 reads, or 98.80% of the total filtered reads.

An analysis of differential abundance of microbiome composition among 12 fecal and 14 gut tissue samples showed significant differential abundance of the bacterial genera, *Apilactobacillus* (Bacillus Firm-5) between queens exposed to 42°C and 32°C, where the log fold change of *Apilactobacillus* significantly increased in queens exposed to heat stress compared to queens in the control (Figure 4, Table S4). We also saw an increase and decrease in the relative abundances of *Bombella* (Alphaproteobacteria) and *Lactobacillus* (Bacillus Firm-4), respectively, in queens exposed to 42°C, however due to the variation in bacterial genera abundance across individual samples (shown in the log-fold-change standard error bars, Figure 4) between heat stress and control queens, differential abundance in other bacterial genera are not statistically significant (Table S4).

**Figure 4.**
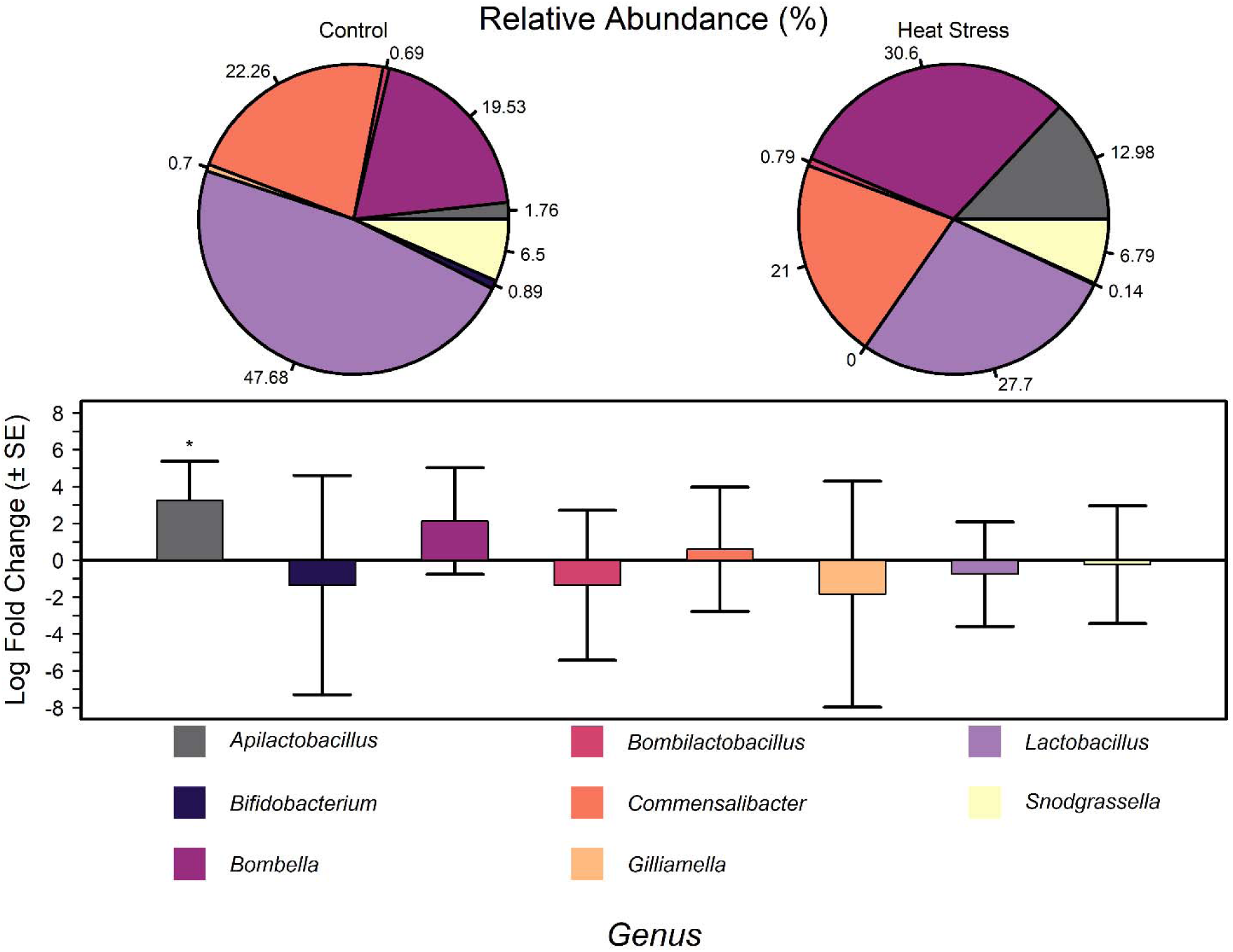
Relative abundance and log-fold-change (± SE) of the bacterial genera among queen honey bees exposed to heat stress or control temperatures for 4 hours (i.e., 42°C or 32°C, respectively). Fecal and gut samples were collected 68h post temperature treatment and sequenced using 16S rRNA amplicon sequencing. Log-fold-changes of bacterial genera abundance is shown for queens exposed to heat stress. Significant differential abundance of bacterial genera among queens exposed to heat stress and control queens are indicated with an asterisk (p < 0.05).

### 4) Refining fecal collection techniques for queen honey bees

Queen fecal deposits were successfully collected using all four techniques, including filter paper, sterile cotton swab, direct transfer using a micropipette, and direct transfer using a micropipette after adding 200µl of molecular water to the fecal deposit. Of these techniques, the direct transfer technique with no water addition was the least preferred, as some deposits were too small to collect with this technique. Based on visual evaluation of DNA bands after gel electrophoresis, of the four sample collection techniques tested and the two DNA extraction kits used (i.e., the Qiagen DNeasy UltraClean microbial and Qiagen QIAamp DNA microbiome kits), the best results were obtained using the microbial kit combined with either the sterile cotton swab or filter paper collection techniques, and the microbiome kit combined with the micropipette transfer technique after molecular water addition. (Figure 5).

**Figure 5.**
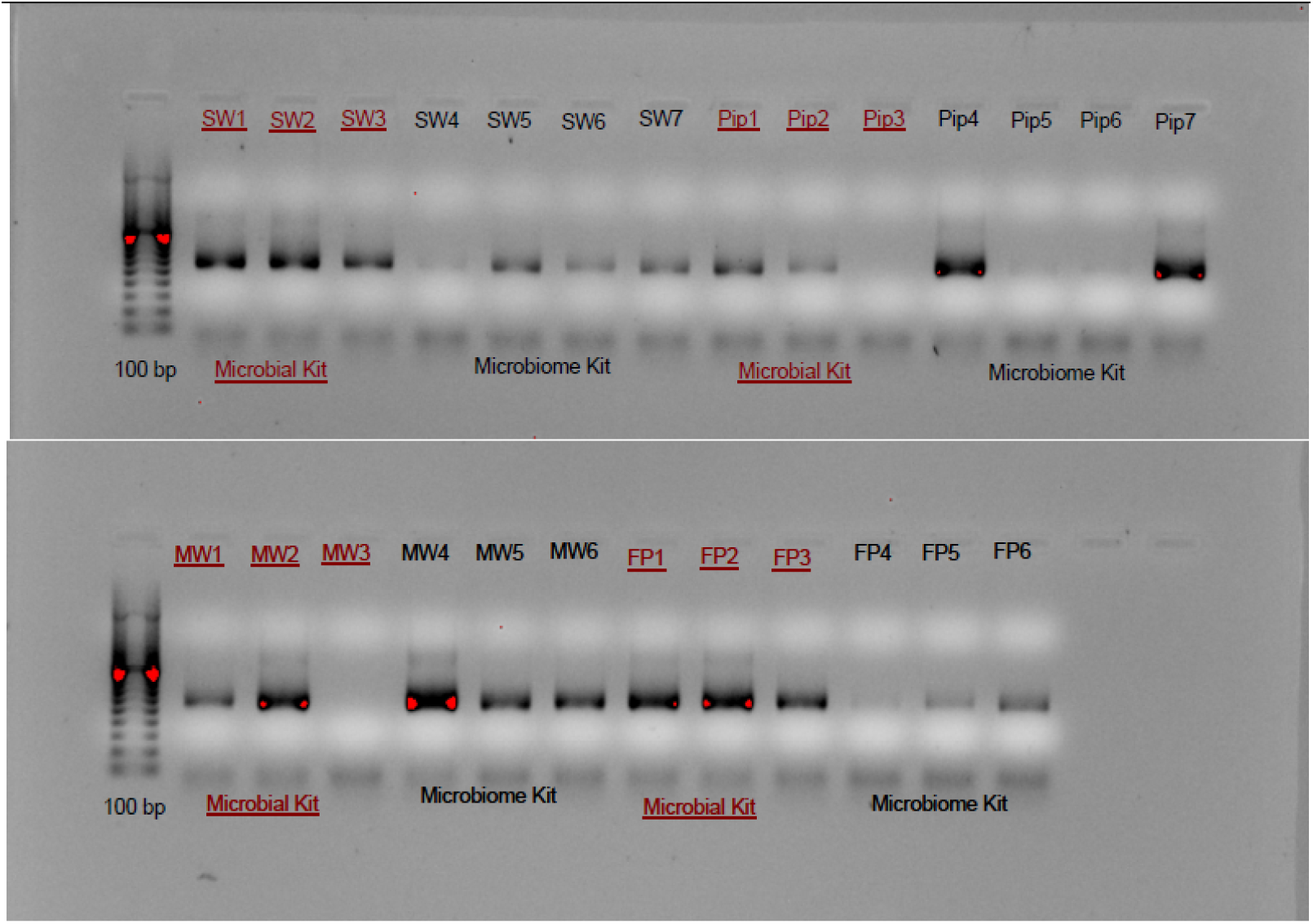
Gel electrophoresis of queen feces showing DNA separation after DNA extraction using the Qiagen DNeasy UltraClean microbial kit (lanes labelled red) or the Qiagen QIAamp DNA microbiome kit (lanes labelled black). Lane one on the top and bottom is the molecular weight ladder. Top: lanes two to eight are fecal samples collected using a sterile cotton swab (i.e., SW1 to SW7), and lanes nine to 15 are fecal samples collected using direct transfer with a micropipette (i.e., Pip1 to Pip7). Bottom: lanes two to seven are fecal samples collected using direct transfer with a micropipette after adding 200µl of molecular water (i.e., MW1 to MW6), and lanes eight to 13 are fecal samples collected using filter paper (i.e., FP1 to FP6).

### 5) Screening for known honey bee viruses in queen honey bee feces

Six of nine honey bee viruses screened were detected in both worker gut and queen fecal samples (Figure 6, Table S5). The three viruses in the screening panel that were not detected in queen feces, were also not detected in the worker gut samples, suggesting CBPV, KBV, and ABPV were either absent from all samples or present below the detection limit (i.e., 10^-10^2^ viral gene copies). Successful virus detection did not differ between worker guts and queen feces (z = −1.18, p = 0.24), however, significantly higher normalized viral gene copy number was observed in queen feces compared to worker guts sampled from the same colony (z = −10.29, p < 0.001, Figure 6, Table S6). In addition, BQCV was detected in significantly higher viral gene copy numbers than all other viruses regardless of sample type (i.e., queen feces or worker guts), but no significant differences in viral gene copy number among the other five viruses were observed (Table S6).

**Figure 6.**
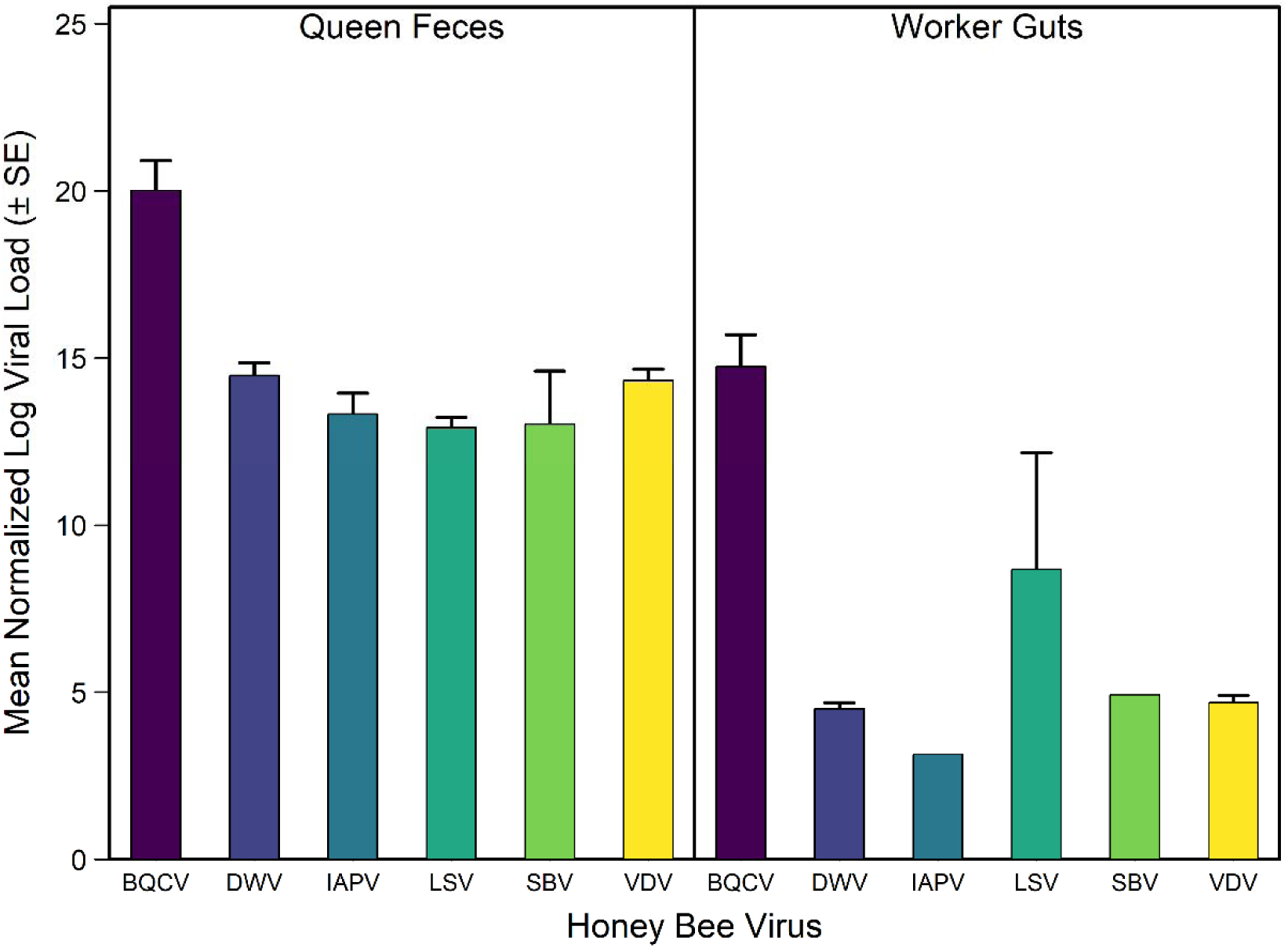
Normalized log viral load (±SE) per queen or worker bee of six viruses (i.e., black queen cell virus (BQCV), deformed wing virus A (DWV), Israeli acute paralysis virus (IAPV), Lake Sinai virus (LSV), sacbrood virus (SBV), and *Varroa destructor* virus (VDV; also known as deformed wing virus B) detected in queen feces (N = 4) and worker guts (N = 4). Worker gut samples were taken from the same colonies (at the same time) as the queens that had their feces sampled. Viral load for workers is per bee, where 15 worker guts were homogenized per sample.

## DISCUSSION

We developed a non-destructive method to characterize the gut microbiome of honey bee queens using fecal sampling. Our optimized protocol enables the reliable collection of at least two fecal deposits per queen within a 6–8-hour window, allowing two investigators to sample 20–25 queens in this period. We found that a minimum of two fecal deposits is required to obtain sufficient 16S rRNA amplicon sequencing data, with no significant increase in sequencing reads when additional fecal deposits are collected from the same queen (Figure 1). This plateau is likely due to a progressive reduction in fecal volume as the queen’s rectum empties and cannot immediately refill. While extending the sampling window could allow queens to replenish their rectal contents, facilitating the collection of larger fecal deposits, the potential variability in the gut microbiome composition between fecal sampling events more than eight hours apart remains unknown.

Over three years, we refined our queen fecal collection methods to improve reliability, efficiency, and downstream DNA extraction and amplification. For optimal sterility and DNA yield, collecting feces deposited in a sterile petri dish using sterile cotton swabs was the preferred approach. Directly transferring the fecal deposit after adding 200 µl of molecular-grade water yielded comparable results but was more time consuming and full sample transfer was rarely achieved. This may explain the poor DNA amplification observed in two of the three samples that were directly transferred after adding molecular-grade water using the Qiagen DNeasy UltraClean microbial kit (Figure 5). The filter paper method also performed as well as the cotton swab and molecular water direct transfer method, but posed a greater risk of contamination, as fecal-stained sections of filter paper require handling during partitioning. The QIAamp DNA microbiome kit performed similarly for samples collected after molecular water application, however, it was less effective than the microbial soil kit collection via a sterile cotton swab or filter paper. Given its significantly lower cost (i.e., the Qiagen QIAamp DNA microbiome kit is approximately five times the cost of the Qiagen DNeasy UltraClean microbial kit), the microbial kit provided a cost-effective alternative without compromising performance.

The bacterial community composition of paired honey bee queen feces and gut tissue samples was similar, supporting the use of queen feces as a non-destructive proxy for gut microbiome characterization. This is reinforced by the significant positive correlations in 16S amplicon sequence reads between fecal and gut samples from individual queens (Figure 2), and the non-significant analysis of similarities (Figure 3). Additionally, all six bacterial genera detected in whole gut tissues were also detected in similar relative abundances in the fecal samples.

Recent studies have cautioned against using feces as a proxy for gut microbiome characterization in other species due to variation in bacterial composition across gut regions (Lam and Fong, 2024; Yan *et al.,* 2019). The honey bee worker gut microbiota similarly varies by gut section (reviewed in Raymann and Moran, 2018; Kwong and Moran, 2016), however, no studies to date have characterized bacterial composition across the entire queen gut. Our findings indicate that whole-gut bacterial communities of queens are well-represented in fecal samples, providing a non-destructive approach for studying the queen honey bee gut microbiome.

Exposure to heat stress altered the bacterial composition and community structure of the queen honey bee gut microbiome, indicating that gut microorganisms respond dynamically to thermal stress (Figure 4). While the thermal tolerance of the queen gut microbiome remains poorly understood, previous work demonstrated that *Snodgrassella* and *Gilliamella* can grow at temperatures exceeding 42°C in culture (Hammer, Le, and Moran, 2021). Consistent with this, we observed minimal changes in the relative abundance of these genera under heat stress. In contrast, *Apilactobacillus* abundance increased, coinciding with a decline in *Lactobacillus*. One possible explanation for this observation is that strains of *Apilactobacillus kunkeei* possess antimicrobial genes encoding for immunity with antimicrobial activity against some *Lactobacillus* commensals in honey bee workers (Zendo *et al*., 2020). Alternatively, heat stress may have directly impacted *Lactobacillus*, as some strains exhibit peak growth rates at 40– 41°C but decline rapidly at higher temperatures (Palmer-Young *et al*., 2023). However, the thermal tolerance of *A. kunkeei* is not well understood, and without species-level resolution in our dataset, the precise relationship between *Apilactobacillus* and *Lactobacillus* under thermal stress requires further investigation.

Our results confirm the successful detection of viruses in queen feces (Figure 6), demonstrating the feasibility of non-destructive virome characterization in honey bee queens. The nine detected viruses are commonly associated with honey bees, including queens, worldwide (Amiri *et al*., 2021; Amiri *et al*., 2020; Chen and Siede, 2007). Honey bee viruses can spread through multiple transmission routes, including venereal, horizonal, vertical, fecal-oral, and vector-borne transmission with *Varroa destructor* acting as a key vector (Chen and Siede, 2007). Interestingly, while we detected six of the nine viruses in both queens and workers (Figure 6), Kevill *et al*., (2020) found that not all viruses present in workers were detectable in queens, suggesting that the worker virome may not always serve as a reliable proxy for queen infection status. Furthermore, although Kevill *et al*., (2020) reported comparable viral loads in queens and workers, viral loads were quantified per sample rather than per individual, with one queen per sample compared to 50 workers. Worker samples may therefore have been overestimated (Kevill *et al*., (2020). Previous studies have not reported viral loads in queen feces that tested positive for viruses (Chen et al., 2006; Hung, 2000). Our non-destructive method enables direct quantification of viral titers in queens, providing new opportunities to investigate viral dynamics in honey bee queens.

The detection of viruses in queen feces offers a non-destructive approach to studying viruses in queens and assessing their health prior to importing, shipping, and colony installation. Although previous studies have detected viruses in queen feces (Chen *et al*., 2006; Hung, 2000), and proposed its use for pre-importation screening (Hung, 2000), this approach has not been widely adopted or even further explored. More recently, Amiri *et al*. (2020) suggested using queen feces to characterize virus infections and assess immune responses in queens, however this was not pursued as fecal sampling was assumed to be challenging. Here, we establish a robust method for collecting queen feces and confirm its utility for virus detection, providing a practical tool for virome surveillance and pre-shipment health assessments in honey bee queens.

Our findings establish proof-of-concept that viruses can be detected in queen honey bee feces and that our fecal collection methods are both reliable and efficient. However, there is still much work to be done to validate the potential of queen feces to characterize and quantify viruses in queen honey bees for the purpose of monitoring and characterizing immune response. Specifically, queens exhibited significantly higher overall viral loads compared to worker guts sampled from the same colonies. The underlying cause of this discrepancy remains unclear—potential explanations include differences in water content between feces and gut homogenates, viral shedding dynamics, lack of correlation between viral loads in queens and workers, or low sample size. Additionally, while expression levels of the housekeeping gene RP49 were consistent within sample types (i.e., feces or worker guts), significant differences were observed between sample types (Table S6), highlighting the need for further investigation into alternative housekeeping genes for normalizing viral gene copy numbers in queen feces. To fully validate the use of queen feces as a tool for virome monitoring, direct comparisons between the queen fecal virome and the queen’s internal virome are necessary, rather than comparisons to worker viromes. However, as a proof-of-concept, we used worker guts as a control, as they are the standard test material for virus detection in honey bee colonies.

Our non-invasive method for characterizing the gut microbiome and virome of queen honey bees, combined with advances in next-generation sequencing, enables, for the first time, longitudinal studies of individual queens throughout their adult lives. This breakthrough provides an unprecedented opportunity to investigate queen health and quality over time.

Repeated sampling of an individual queen’s gut microbiome and virome allows us to explore key questions about the relationships between microbiome composition, immune function (Mason and Shikano, 2023; Sepulveda and Moeller, 2020), and reproduction (Zhang et al., 2019). Given the demonstrated trade-off between stored sperm viability and immune function in queen honey bees (McAfee et al., 2021), our method enables targeted investigations into how temperature-induced gut dysbiosis impacts queen health and fertility. Beyond temperature stress, these methods can be extended to assess the effects of other stressors, including antibiotic and pesticide exposure, providing insights into the long-term consequences of environmental challenges on queen performance and resilience. Finally, regulatory agencies assess pathogen transmission in their risk assessments of importing and exporting honey bees (Canadian Food Inspection Agency, 2024; World Organisation for Animal Health (OIE), 2010; Code of Federal Regulations, 2004), considering potential impacts on other honey bees, livestock, and native or non-native species. Our non-destructive approach to evaluating queen health offers a practical tool to facilitate queen trade, enhance biosecurity measures, and assess the viability of queens transported across regions and borders.

## ONLINE METHODS

### Sampling queen feces and gut tissues

Feces and gut tissues from 47 locally raised queens (less than one year old) in Lethbridge, Alberta, Canada, were sampled in 2022. Queen bees were removed from their colonies, caged with five attendants in four-hole-wooden cages (i.e., a small block of wood with four connecting wells covered in a mesh screen), provided queen candy (i.e., mixture of confectioners’ sugar and honey), and brought into the lab. To sample feces, each queen was removed from her respective cage and placed into a sterile plastic 50 mm petri dish with a sterile 47 mm 0.2 µm nitrocellulose membrane filter paper disc (Merck Millipore Ltd., Ireland). Each queen was then continuously observed for defecation behaviour. Immediately upon defecation, each queen was placed into a new sterile 50 mm petri dish arena with a new sterile filter paper disc to collect any additional fecal deposits prior to dissection of the queen. All filter paper discs with feces were placed into sterile 5 ml plastic tubes and immediately stored in a −80°C freezer.

Queens were euthanized by decapitation and their entire gut (e.g., midgut, ileum, and rectum) was removed by holding the thorax of the queen and gently pulling on the stinger with sterile forceps. The entire gut was placed on a sterile 90 mm plastic petri dish on ice, the stinger was removed, the gut placed in a sterile 1.5 ml microcentrifuge tube and immediately stored at - 80°C. All dissection tools were sterilized between queens with 90% ethanol and new sterile petri dishes were used for each dissection.

At the start of the project, we were uncertain of the number of fecal deposits needed to successfully obtain sequencing data. Thus, working with 47 queens, we pooled one to four fecal deposits to create pooled samples and separately pooled one to four guts per samples (Table 1). As a single queen would often provide more than one fecal deposit, the number of fecal deposits tended to be larger than the number of queen guts pooled from the same queens (Table 1). After pooling, fecal and gut samples from the 47 queens comprised 21 fecal and 21 gut samples; these samples were used to determine how many fecal deposits are required for sequencing (objective 1), whether the fecal microbiome is representative of the gut microbiome in queen honey bees (objective 2), and a subset of queens was used to characterize the fecal and gut microbiome of queens in response to temperature stress (objective 3).

#### 1) Determining the number of queen fecal deposits needed for successful 16S rRNA amplicon sequencing

Feces from the 47 queens were pooled into 21 fecal samples as described above. We tested sequencing of one to four fecal deposits per sample (Table 1) to determine the minimum number of queen fecal deposits required to obtain sequencing data.

#### 2) Characterizing the fecal and gut microbiome communities of queen honey bees

To determine whether the fecal microbiome is correlated with the gut microbiome in queen honey bees, the feces and guts collected from 47 queens and pooled into 21 fecal and 21 gut samples (as described above)(Table 1) were used. Of these, 19 fecal samples (and their paired gut samples) were used as they passed quality control (i.e., >=1000 reads).

#### 3) Fecal and gut microbiome communities of queen honey bees in response to temperature stress

To test whether queen feces can be used to screen for possible temperature stress exposure, thirty-one of the queen bees were divided between two temperature treatments. Small wooden cages containing a single queen bee, and five attendant worker bees were exposed to either 42°C or 32°C for four hours and then subsequently kept in a growth chamber set to 32°C for 68 hours prior to fecal sampling and dissection. Queens and attendants were provided queen candy and water several times a day throughout the 68-hour holding period. We chose a 68-hour holding period to allow time for the microbiome to respond and ensure changes to gut microbiome in response to temperature would be reflected in the gut and fecal samples, as three days was the minimum time for Raymann, Shaffer, and Moran (2017) to observe changes to the worker honey bee gut microbiota after antibiotic exposure. Fecal and gut sampling and pooling is described above (Table 1). The same two fecal samples with a low number of reads that were removed from objective two analyses, were also removed from this analysis, but their paired gut samples were included to increase sample size of the temperature treatment data since correlations between fecal and gut samples was not of interest for this objective. Thus, we obtained sequencing data from 12 pooled fecal and 14 pooled gut samples.

### DNA Extraction, Amplification, Library Prep, and 16S Amplicon Sequencing (Objectives 1-3)

Twenty-one fecal and 21 gut samples pooled (Table 1) from the 47 queens were sent to Microbiome Insights (Richmond, British Columbia, Canada) for DNA extraction, amplification, library prep, and 16S rRNA amplicon sequencing. DNA was extracted using the Qiagen MagAttract PowerSoil DNA KF kit (Formerly MO BioPowerSoil DNA Kit) using a KingFisher robot. DNA quality was visually evaluated on gel electrophoresis and quantified on a Qubit 3.0 fluorometer (Thermo-Fisher, Waltham, MA, USA). Library prep was completed with the Illumina Nextera kit (Illumina, San Diego, CA, USA).

Bacterial 16S rRNA genes were amplified with dual-barcoded primers targeting the V4 region following Kozich *et al*. (2013). Amplicons were sequenced with an Illumina MiSeq using the 300-bp paired end kit (v.3). Sequences were denoised, taxonomically classified using the reference database Greengenes (v. 13_8), and clustered into 97% similarity operational taxonomic units (OTUs) with the *mothur* software package (v. 1.39.5) (Schloss *et al.,* 2009).

The potential for contamination was addressed by co-sequencing DNA amplified from our samples and from template-free controls and extraction kit reagents. Two positive controls, consisting of cloned SUP05 DNA, were also included (number of copies = 2*10^6). Operational taxonomic units were considered putative contaminants and removed if their mean abundance in controls were 25% or higher of their mean abundance in our samples.

### Microbiome Analysis (Objectives 1-3)

Illumina sequence reads were processed in the R software environment version 4.4.1 (R Core Team, 2023) following the dada2 pipeline version 1.16 (https://benjjneb.github.io/dada2/tutorial.html) (Callahan *et al.,* 2016). FASTQ files were filtered and trimmed with a truncLen = c(240, 40) and a maxEE = c(2,5) to account for poor-quality reverse reads. Forward and reverse Illumina reads were joined, and chimeras were removed. Taxonomy was assigned to amplicon sequence variants (ASVs) using BEExact v2023.01.30 training set DADA2 V4 (Daisley and Reid, 2021) and the Silva 138.1 prokaryotic SSU taxonomic training set with species assignment for DADA2 (Callahan *et al.,* 2016; Yilmaz *et al.,* 2014; Quast *et al.,* 2013; Pruesse *et al.,* 2007). For objectives two and three, two fecal samples with less than 1000 sequence reads were removed from the dataset along with their paired pooled gut samples (where appropriate (i.e., objective 2) prior to FilterandTrim (DADA2), such that error rates were learned from the filtered FASTQ sequence files included in the sample analyses. For this reason, the number of sequenced reads before rarefaction and downstream analyses varies by objective and are thus reported for each objective in the results.

#### 4) Refining fecal collections from queen honey bees

In 2024, locally raised queens (less than one year old) were used to further refine our fecal collection methods. Twenty-six queens were collected from their colonies, caged with five attendant workers in new four-hole-wooden cages, and brought into the lab for fecal sampling. The previous method of using full discs of filter paper to collect queen feces was problematic for DNA extractions because it was difficult to homogenize the filter paper in a small volume of buffer. Therefore, we explored other methods of collecting queen feces from the petri dish arenas. We compared a slightly modified version of our filter paper technique to three other sampling techniques, including the use of a sterile cotton swab, direct transfer using a micropipette, and direct transfer using a micropipette after adding 200 µl of molecular water (i.e., DNAse/RNAse free water) to the fecal deposit. Queens assigned to the filter paper technique were placed into a sterile 50 mm petri dish with a sterile 47 mm 0.2 µm nitrocellulose membrane filter paper disc (Merck Millipore Ltd., Ireland), while queens assigned to the other three sampling techniques were placed directly into a sterile 50 mm petri dish. Queens were continuously observed for defecation and immediately removed from their arenas upon defecation.

To reliably collect more than one fecal deposit from each queen, queens were returned to their respective cages with attendants, provided water and queen candy, and stored in a dark drawer at room temperature for 3-4 h. Queens were once again removed from their cages and placed into a new sterile arena either with or without a filter paper disc, (i.e., depending on fecal collection treatment technique) and an additional fecal deposit was collected.

Our filter paper technique was slightly modified whereby the section of the filter paper that contained the fecal deposit was cut from the whole disc and placed into a sterile 1.5 ml microcentrifuge tube containing 200 µl of molecular water. For the cotton swab technique, a sterile cotton swab was used to soak up the fecal deposit from the Petrie dish and the tip of the swab was cut into a sterile 1.5 ml microcentrifuge tube containing 200 µl of molecular water.

Filter paper discs and cotton swab tips were cut using micro dissecting scissors sterilized in 90% ethanol. For the direct transfer methods without molecular water, micropipettes were used to transfer the fecal deposit directly from the surface of the petri dish arena to a sterile 1.5 ml microcentrifuge tube containing 200 µl of molecular water. For the direct transfer method with molecular water, 200 µl of molecular water was added to the fecal deposit before directly transferring the volume of molecular water comprising the fecal deposit from the surface of the petri dish to a sterile 1.5ml microcentrifuge tube. Samples were immediately stored in a −80°C freezer until they were shipped to the National Bee Diagnostic Center (NBDC) in Beaverlodge, Alberta, Canada for DNA extraction and amplification. Each of the four sampling techniques was replicated six or seven times, with each fecal sample including at least two separate fecal deposits from the same queen.

### DNA Extraction and Amplification (Objective 4)

Queen feces collected via filter paper discs and cotton swabs were extracted by transferring the contents of the 1.5 ml microcentrifuge tubes (i.e., fragments of filter paper or fragments of cotton swab and 200 µl of molecular water to 2 ml screw cap tubes with 2.8 mm beads. 400 µl of nuclease-free water was added to each tube and homogenized in a bead beater for 30 s at 6 m/s. Homogenized samples were centrifuged at 5000 rpm for 30 s and then 20 s and the supernatant was transferred to a 1.5 ml tube at the end of each cycle. The supernatant was centrifuged at 10,000 rpm for 3 min to precipitate a pellet.

Queen feces collected via the direct transfer with and without molecular water methods did not require transferring or pre-processing. We tested two DNA extraction kits for each fecal sample collection technique, including the Qiagen QIAamp DNA Microbiome Kit protocol (May 2014) and the Qiagen DNeasy UltraClean Microbial Kit, following the kit protocols. The six or seven fecal deposits collected for each sampling technique were divided among the two DNA extraction kits, such that three or four samples for each fecal collection technique was assigned to each DNA extraction kit.

Bacterial 16S rRNA genes were amplified with dual-barcoded primers targeting the V3 and V4 regions, using forward primers (5’-TCGTCGGCAGCGTCAGATGTGTATAAGAGACAGCCTACGGGNGGCWGCAG) and reverse primers (5’-GTCTCGTGGGCTCGGAGATGTGTATAAGAGACAGGACTACHVGGGTATCTAATCC). DNA quality was visually evaluated on gel electrophoresis to determine which fecal collection technique and DNA extraction kit was optimal for sampling queen honey bee feces.

#### 5) Screening for known honey bee viruses in queen honey bee feces

Additional locally raised mated queens (0-2 months old) were used to screen for honey bee viruses in queens in 2024. Four queens were collected from their colonies in Lethbridge, Alberta and caged in new four-hole-wooden cages with five attendant workers from the same colony. After caging each queen, a 50 ml plastic tube was used to sample approximately 100 worker bees from the same frame the queen was located. Worker samples were immediately placed in a cooler of ice, and caged queens were returned to their respective colonies for 16-20 hours prior to fecal sampling. We established over the three years of developing these methods, that this holding period in the field ensures the queen has a full rectum for efficient fecal sampling the next day. The next morning, the caged queens were brought into the lab for fecal sampling. Fecal sampling followed the same methods described in our refined methods above using sterile cotton swabs.

Screening for viruses in honey bee colonies is typically done on a sample of adult workers, where workers are homogenized before analysis. Thus, fifteen workers from each of the four paired worker samples (i.e., workers sampled from the same colony frame their queens were collected from) were dissected by holding the thorax of each worker and gently pulling on the stinger with sterile forceps (Tarpy, Mattila, and Newton, 2015) and their entire gut (e.g., midgut, ileum, and rectum) was isolated and homogenized in one sterile 1.5 ml microcentrifuge tube.

The 1.5 ml microcentrifuge tubes of homogenized worker guts were stored on ice until all 15 workers in each sample were dissected. Worker gut samples were then stored in a −80°C freezer. Queen fecal samples (n = 4) and their paired pooled worker gut samples (n = 4) were then shipped to the NBDC for viral analysis.

The samples were screened for nine viruses commonly found in honey bees: Israeli acute paralysis virus (IAPV), deformed wing virus A (DWV-A), *Varroa destructor* virus (VDV; also known as deformed wing virus B or DWV-B), acute bee paralysis virus (ABPV), Kashmir bee virus (KBV), chronic bee paralysis virus (CBPV), black queen cell virus (BQCV), Lake Sinai virus (LSV), and sacbrood virus (SBV). Worker guts and queen fecal samples were homogenized in 400 µl and 200 µl of GITC buffer, respectively (Evans *et al*., 2013). An aliquot of 200 µl was used to isolate total RNA using the NucleoSpin®RNA kit following manufacturer instructions (Macherey-Nagel Gmbh & Co. KG, Düren, Germany). RNA was quantified using IMPLEN Nanophtometer NP80 (Munich, Germany). cDNA was synthesized from 800 ng of total RNA for 20 minutes at 46°C in a final volume of 20 µl using the iScript cDNA synthesis kit (Bio-Rad laboratories, Hercules, USA). cDNA was diluted with 60 µl of molecular grade water to a total of 80 µl from which 3 µl (30 ng) were used for qPCR quantification.

Quantification of viral infection levels was determined by real-time PCR using primers (Table S1) and SSoAdvanced™ Universal SYBR® Green Supermix (Bio-Rad Laboratories, Hercules, USA). Amplification assays were performed by triplicate employing ∼30 ng of cDNA (Viruses) in a CFX384 Touch™ Real-Time Detection System (Bio-Rad Laboratories, Hercules, USA).

RP49 was chosen as reference gene. Standard curves were prepared from plasmids harboring the target amplicons with copy numbers diluted from 10^7^ to 10^2^. PCR conditions were 3 min at 95°C for initial denaturation/enzyme activation followed by 40 cycles of 10 s at 95°C and 30 s at 60°C (except IAPV, where annealing/extension was 45 s at 60°C). Specificity was checked by performing a melt-curve analysis 65-95°C with increments of 0.5°C 2 s/step. Results were analyzed with the CFX Manager™ Software and exported to an Excel spreadsheet to calculate virus copy numbers per sample. Virus copy numbers per bee were calculated after the viral data were exported from the BioRad CFX Manager™ Software, where worker samples had 15 workers per sample and queen fecal samples had one queen per sample.

### Statistical Analysis

All statistical analyses were performed in the R software environment version 4.4.1 (R Core Team, 2023).

#### 1) Determining the number of queen fecal samples needed for successful 16S rRNA amplicon sequencing

To determine the minimum number of fecal deposits necessary to successfully sequence 16S amplicon reads from queen honey bee feces, sequence data from fecal samples (n = 21) were filtered to remove ASVs where zero reads were sequenced across the samples because for this objective, we were interested in the number of reads sequenced per sample and not the representation of specific taxonomic ASVs sequenced in a sample. A generalized linear model (GLM) with a negative binomial link error distribution, using the *mgcv* package (Wood 2004) was used to characterize mean number of 16S amplicon sequence reads across the categorical variable number of pooled fecal deposits (i.e., 1, 2, 3, or 4 (Table 1)). Statistical significance was evaluated using model selection with Akaike Information Criteria (AIC) (Burnham and Anderson 2002, Wood et al. 2016, Wood 2017). The general linear hypothesis testing function *glht* in the *Multcomp* package (Hothorn et al. 2008) was used for post-hoc analysis of GLMs; p values were adjusted for Type I error using the Bonferroni method.

#### 2) Characterizing the fecal and gut microbiome communities of queen honey bees

To determine whether the microbiome found in queen feces is representative of the microbiome found in queen gut tissues, we examined the linear relationship between the log abundance of 16S amplicon sequence reads found in the paired 19 fecal and 19 gut tissue samples and performed Pearson correlation analyses for each bacterial genera represented using the *cor.test* function (R Core Team, 2023). We also performed an analysis of similarities (ANOSIM) on a pairwise Bray-Curtis dissimilarity matrix using the *anosim* function in the vegan package (Oksanen *et al.,* 2022) to test for significant differences in bacterial community composition among the paired 19 fecal and 19 gut samples.

#### 3) Fecal and gut microbiome communities of queen honey bees in response to temperature stress

To determine whether the microbiome of queen honey bees exposed to heat stress (i.e., 42°C) significantly differed from queens exposed to control temperatures (i.e., 32°C), we performed an analysis of differential abundance using the *aldex.clr* function in the *ALDEx2* package (Fernandes *et al*., 2013; Gloor, Macklaim, and Fernandes, 2016; Fernandes *et al*., 2022). The analysis was performed to identify bacterial taxa that vary in abundance between queens exposed to heat stress and control queens and used 1000 Monte Carlo instances to estimate the underlying Dirichlet distributions due to a small number of taxa sequenced in queen honey bee feces and guts. A Wilcoxon Rank Sum test was used to test for significant differences among the centred-log-ratio transformed taxon abundance with p-value adjustments using Benjamini-Hoschberg corrections using the *aldex.ttest* function in the *ALDEx2* package (Fernandes *et al*., 2013; Gloor, Macklaim, and Fernandes, 2016; Fernandes *et al*., 2022).

#### 5) Screening for known honey bee viruses in queen honey bee feces

A GLM with a binomial link error distribution, using the *mgcv* package (Wood 2004) was used to characterize the detection of honey bee viruses across the categorical variable, sample type (i.e., queen feces and worker guts) for each virus screened (i.e., DWV, CBPV, VDV, SBV, BQCV, IAPV, KBV, LSV, and ABPV). Statistical significance was evaluated using model selection with AIC (Burnham and Anderson 2002, Wood et al. 2016, Wood 2017). The general linear hypothesis testing function *glht* in the *Multcomp* package (Hothorn et al. 2008) was used for post-hoc analysis of GLMs; p values were adjusted for Type I error using the Bonferroni method. A GLM with a gaussian link error distribution and model selection with post-hoc analysis was also used to characterize the log-transformed viral load per bee (i.e., the normalized gene copies of each virus) across the categorical variables sample type (i.e., queen feces and worker guts) and viruses (i.e., DWV, CBPV, VDV, SBV, BQCV, IAPV, KBV, LSV, and ABPV).

## ACKNOWLEDGEMENTS

We sincerely thank Jeff Kearns, Kelsey Gourlie, Jessenia Buzunis-Delagneau, Jemma Todoschuk, Donaya Baker, and Fairo Dzekashu Foryuy for their help in managing honey bee colonies, collecting queens from the field, and queen fecal sampling. Funding for this project was provided by Project Apis m, MITACS, RDAR, the Alberta Beekeepers Commission, and the Government of Canada through the Agriculture and Agri-Food Canada Genomics Research and Development Initiative.

## AUTHOR CONTRIBUTIONS

LAH, SEH, JP, and MMG conceived and planned the experiments and acquired funding. LAH, PWV, and SEH carried out the experiments and analyzed samples. LAH led the data analysis and LAH and SEH led the original draft writing. All authors contributed critically to the final draft and gave final approval for publication. All authors declare no conflicting interests.

## DATA AVAILABILITY

The data used for this study is available from the corresponding authors on reasonable request.

