## Supplementary material for "A non-invasive method for profiling the gut microbiome and virome of honey bee queens": Table S

**Table S1.** Primers, their sequence, and amplicon size in base pairs (bp) used to detect and quantify honey bee viruses sampled from queen honey bee feces and worker honey bee guts (objective 5). Viruses included Acute Bee Paralysis virus (ABPV), Kashmir Bee Virus (KBV), Chronic Bee Paralysis virus (CBPV), Lake Sinai virus (LSV), deformed wing virus (DWV), *Varroa destructor* virus (VDV), Black queen cell virus (BQCV), Israeli acute paralysis virus (IAPV), and Sacbrood virus (SBV). RP49 is the ribosomal protein 49.

| Target | Primer | Sequence (5'-3') | Amplicon size (bp) | Reference |
|---|---|---|---|---|
| ABPV | ABPV-F6548 | TCATACCTGCCGATCAAG | 197 | Locke *et al*., 2012 |
| | KIABPV-B6707 | CTGAATAATACTGTGCGTATC | | |
| KBV | KBV-F | TGAACGTCGACCTATTGAAAAA | 106 | VanEnglesdorp *et al*., 2009 |
| | KBV-R | TCGATTTTCCATCAAATGAGC | | |
| CBPV | CBPV1-qF1818 | CAACCTGCCTCAACACAG | 296 | Locke *et al*., 2012 |
| | CBPV1-qB2077 | AATCTGGCAAGGTTGACTGG | | |
| LSV 1-4 | LSV1-4-F-2157 | CGTGCGGACCTCATTTCTTCATGT | 152 | Daughenbaugh *et al*., 2015 |
| | LSV1-4-R-2309 | CTGCGAAGCACTAAAGCGTT | | |
| DWV | DWV-F8668 | TTCATTAAAGCCACCTGGAACATC | 136 | Locke *et al*., 2012 |
| | DWV-B8757 | TTTCCTCATTAACTGTGTCGTTGA | | |
| VDV | VDV-F2 | TATCTTCATTAAAACCGCCAGGCT | 140 | McMahon *et al*., 2015 |
| | VDV-R2a | CTTCCTCATTAACTGAGTTGTTGTC | | |
| BQCV | BQCV-qF7893 | AGTGGCGGAGATGTATGC | 294 | Locke *et al*., 2012 |
| | BQCV-qB8150 | GGAGGTGAAGTGGCTATATC | | |
| IAPV | IAPV-F1aF | GCGGAGAATATAAGGCTCAG | 587 | VanEnglesdorp *et al*., 2009 |
| | IAPV-F1a R | CTTGCAAGATAAGAAAGGGGG | | |
| SBV | SBV-qF3164 | TTGGAACTACGCATTCTCTG | 335 | Locke *et al*., 2012 |
| | SBV-qB3461 | GCTCTAACCTCGCATCAAC | | |
| RP49 | RP49-qF | AAGTTCATTCGTCACCAGAG | 205 | Locke *et al*., 2012 |
| | RP49-qB | CTTCCAGTTCCTTGACATTATG | | |

Daughenbaugh, K. F., Martin, M., Brutscher, L. M., Cavigli, I., Garcia, E., Lavin, M., & Flenniken, M. L. (2015). Honey bee infecting Lake Sinai viruses. *Viruses, 7*(6), 3285–3309. https://doi.org/10.3390/v7062772

Evans, J. D., Schwarz, R. S., Chen, Y. P., Budge, G., Cornman, R. S., De la Rua, P., de Miranda, J. R., Foret, S., Foster, L., Gauthier, L., Genersch, E., Gisder, S., Jarosch, A., Kucharski, R., Lopez, D., Lun, C. M., Moritz, R. F. A., Maleszka, R., Muñoz, I., & Pinto, M. A. (2013). Standard methods for molecular research in *Apis mellifera*. *Journal of Apicultural Research, 52*(1), 1–54. https://doi.org/10.3896/IBRA.1.52.4.11

Locke, B., Forsgren, E., Fries, I., & de Miranda, J. R. (2012). Acaricide treatment affects viral dynamics in *Varroa destructor*-infested honey bee colonies via both host physiology and mite control. *Applied and Environmental Microbiology, 78*(1), 227–235. https://doi.org/10.1128/AEM.06094-11

McMahon, D. P., Fürst, M. A., Caspar, J., Theodorou, P., Brown, M. J. F., & Paxton, R. J. (2015). A sting in the spit: Widespread cross-infection of multiple RNA viruses across wild and managed bees. *Journal of Animal Ecology, 84*(3), 615–624. https://doi.org/10.1111/1365-2656.12345

VanEngelsdorp, D., Evans, J. D., Saegerman, C., Mullin, C., Haubruge, E., Nguyen, B. K., … & Pettis, J. S. (2009). Colony collapse disorder: A descriptive study. *PLoS ONE, 4*(8), e6481. https://doi.org/10.1371/journal.pone.0006481

Table S2. Model selection using Akaike information criterion (AIC) for the effect of number of pooled fecal deposits (X) (i.e., 1, 2, 3, or 4) on the mean number of 16S amplicon sequence reads, Y, sequenced across 21 samples of pooled fecal deposits (Table 1). A negative binomial distribution was fit to 199 observations with 304.15 null deviance. Data were rarefied to a sequencing depth of 1500 reads. The table also shows the general linear hypothesis post-hoc testing on the top model fit selected by AIC for the effect of number of fecal deposits on the mean number of 16S amplicon reads with adjusted p values for Type I error using the Bonferroni method.

| Model | ΔqAIC | Df | Weight | Residual Deviance |
|---|---|---|---|---|
| Y ~ X | 0.00 | 5 | 1 | 270.50 |
| Y ~ 1 | 25.7 | 2 | 0 | 304.15 |
| **Post-hoc Comparisons** | | **Std. Error** | **z value** | **Pr(>\|z\|)** |
| 1 fecal deposit – 2 fecal deposits | | 0.34 | 4.49 | < 0.001 |
| 1 fecal deposit – 3 fecal deposits | | 0.38 | 5.03 | < 0.001 |
| 1 fecal deposit – 4 fecal deposits | | 0.33 | 5.09 | < 0.001 |
| 2 fecal deposits – 3 fecal deposits | | 0.41 | 0.93 | 1 |
| 2 fecal deposits – 4 fecal deposits | | 0.36 | 0.41 | 1 |
| 3 fecal deposits – 4 fecal deposits | | 0.40 | -0.58 | 1 |

**Table S3.** Pearson correlation analyses of the log abundance of 16S amplicon sequence reads found in 19 queen honey bee fecal and 19 queen gut tissue samples for each bacterial genera (and their class) represented. Queens providing the fecal deposits in a fecal sample are the same queens as the gut samples.

| Bacterial *Genera(s)* | Correlation Coefficient | p – value | 95% Confidence Interval |
|---|---|---|---|
| *Apilactobacillus* (Bacilli Firm-5) | 0.87 | < 0.001 | 0.68, 0.95 |
| *Bifidobacterium* (Actinobacteria) | 0.75 | < 0.001 | 0.45, 0.90 |
| *Bombella* (Alphaproteobacteria) | 0.80 | < 0.001 | 0.54, 0.92 |
| *Bombilactobacillus* and *Lactobacillus* (Bacilli Firm-4) | 0.83 | < 0.001 | 0.70, 0.91 |
| *Commensalibacter* (Alphaproteobacteria) | 0.70 | < 0.001 | 0.36, 0.88 |
| *Snodgrassella* (Gammaproteobacteria) | 0.81 | < 0.001 | 0.57, 0.92 |

**Table S4.** Analysis of differential abundance of bacterial genus composition from 19 fecal and 21 gut samples from queen honey bees exposed to either heat stress (i.e., 42°C) or control (i.e., 32°C) temperatures for 4h. Effect sizes and expected Benjamini-Hochberg corrected p-values of the Wilcoxon Rank Sum tests for each bacterial genus comparison between temperature treatments are reported after computing Monte Carlo samples of the Dirichlet distribution using a centred log-ratio transformation.

| Bacterial *Genera(s)* | Effect Size | Adjusted p – value | 95% Confidence Interval |
|---|---|---|---|
| *Apilactobacillus* (Bacilli Firm-5) | 1.53 | 0.004 | -0.10, 3.68 |
| *Bifidobacterium* (Actinobacteria) | -0.23 | 0.694 | -2.26, 1.40 |
| *Bombella* (Alphaproteobacteria) | 0.75 | 0.440 | -1.03, 3.43 |
| *Bombilactobacillus* (Bacilli Firm-4) | -0.32 | 0.999 | -1.52, 2.55 |
| *Commensalibacter* (Alphaproteobacteria) | 0.18 | 0.100 | -2.41, 1.78 |
| *Gilliamella* (Gammaproteobacteria) | -0.29 | 0.849 | -2.96, 1.41 |
| *Lactobacillus* (Bacilli Firm-4) | -0.26 | 0.772 | -1.63, 1.41 |
| *Snodgrassella* (Gammaproteobacteria) | -0.08 | 0.998 | -2.28, 2.42 |

**Table S5.** Normalized viral gene copies for nine screened honey bee viruses in honey bee worker guts and queen honey bee feces. Viruses included Acute Bee Paralysis virus (ABPV), Kashmir Bee Virus (KBV), Chronic Bee Paralysis virus (CBPV), Lake Sinai virus (LSV), deformed wing virus (DWV), *Varroa destructor* virus (VDV), Black queen cell virus (BQCV), Israeli acute paralysis virus (IAPV), and Sacbrood virus (SBV). Worker gut samples (N = 4) were a composite sample of 15 honey bee worker guts (i.e., midgut, ileum, and rectum). Queen fecal samples (N = 4) were a composite sample of at least two fecal deposits from individual queens. Queens and workers were paired samples, such that the worker samples were of workers taken from the same colony as the respective queen samples. Effect sizes and expected Benjamini-Hochberg corrected p-values of the Wilcoxon Rank Sum tests for each bacterial genus comparison between temperature treatments are reported after computing Monte Carlo samples of the Dirichlet distribution using a centred log-ratio transformation.

| Honey Bee Virus | Normalized Log Virus Gene Copy Number Per Bee | | | | | | | |
|---|---|---|---|---|---|---|---|---|
| | Worker Guts | | | | Queen Feces | | | |
| DWV-A | 2.08 | 1.78 | 2.08 | 1.90 | 6.15 | 6.78 | 6.20 | 6.02 |
| CBPV | ND | ND | ND | ND | ND | ND | ND | ND |
| VDV | 2.26 | 1.89 | 2.12 | 1.87 | 6.00 | 6.21 | 6.04 | 6.65 |
| SBV | 2.14 | ND | ND | ND | ND | 4.96 | ND | 6.35 |
| BQCV | 6.95 | 6.76 | 6.74 | 5.17 | 7.60 | 9.14 | 8.68 | 9.34 |
| IAPV | ND | 1.36 | ND | ND | 5.23 | 5.93 | 6.48 | 5.50 |
| KBV | ND | ND | ND | ND | ND | ND | ND | ND |
| LSV | 1.11 | 3.84 | 3.36 | ND | 5.47 | 5.31 | 5.85 | 5.81 |
| ACBV | ND | ND | ND | ND | ND | ND | ND | ND |

* ND = Not Detected

**Table S6.** Model selection using Akaike information criterion (AIC) for the effect of sample type (X) (i.e., queen feces and worker guts) and virus (V) (i.e., Acute Bee Paralysis virus (ABPV), Kashmir Bee Virus (KBV), Chronic Bee Paralysis virus (CBPV), Lake Sinai virus (LSV), deformed wing virus (DWV), *Varroa destructor* virus (VDV), Black queen cell virus (BQCV), Israeli acute paralysis virus (IAPV), and Sacbrood virus (SBV)) on the mean normalized log viral load detected (Y). A gaussian distribution was fit to 39 observations with 1050.5 null deviance. This table also shows the general linear hypothesis post-hoc testing on the top model fit selected by AIC for the effect of sample type (i.e., queen feces and worker guts) on the mean normalized log viral load of six honey bee viruses and the effect of virus on the mean normalized log viral load detected in queen fecal and worker gut samples with adjusted p values for Type I error using the Bonferroni method.

| Model | ΔqAIC | Df | Weight | Residual Deviance |
|---|---|---|---|---|
| Y ~ X + V | 0.00 | 8 | 0.87 | 163.92 |
| Y ~ X * V | 3.73 | 13 | 0.13 | 108.67 |
| Y ~ X | 33.40 | 3 | 0.00 | 554.31 |
| Y ~ V | 53.77 | 7 | 0.00 | 706.14 |
| Y ~ 1 | 55.97 | 2 | 0.00 | 1050.2 |
| **Post-hoc Comparisons** | | **Std. Error** | **z value** | **Pr(>\|z\|)** |
| Queen feces – Worker guts | | 0.75 | -10.29 | < 0.001 |
| BQCV – DWV | | 1.13 | -6.97 | < 0.001 |
| BQCV – IAPV | | 1.31 | -6.41 | < 0.001 |
| BQCV – LSV | | 1.17 | -5.82 | < 0.001 |
| BQCV – SBV | | 1.54 | -5.42 | < 0.001 |
| BQCV – VDV | | 1.13 | -6.95 | < 0.001 |
| DWV – IAPV | | 1.31 | -0.39 | 1 |
| DWV – LSV | | 1.17 | 0.91 | 1 |
| DWV – SBV | | 1.54 | -0.29 | 1 |
| DWV – VDV | | 1.13 | 0.02 | 1 |
| IAPV – LSV | | 1.34 | 1.18 | 1 |
| IAPV – SBV | | 1.66 | 0.04 | 1 |
| IAPV – VDV | | 1.31 | 0.40 | 1 |
| LSV – SBV | | 1.56 | -0.97 | 1 |
| LSV – VDV | | 1.17 | -0.89 | 1 |
| SBV – VDV | | 1.54 | 0.30 | 1 |
